# The dynamics of motor learning through the formation of internal models

**DOI:** 10.1101/652727

**Authors:** Camilla Pierella, Maura Casadio, Sara A. Solla, Ferinando A. Mussa-Ivaldi

## Abstract

A medical student learning to perform a laparoscopic procedure as well as a recently paralyzed user of a powered wheelchair must learn to operate machinery via interfaces that translate their actions into commands for the external device. Mathematically, we describe this type of learning as a deterministic dynamical process, whose state is the evolving forward and inverse internal models of the interface. The forward model predicts the outcomes of actions while the inverse model generates actions designed to attain desired outcomes. Both the mathematical analysis of learning dynamics and the performance observed in a group of subjects demonstrate first-order exponential convergence of the learning process toward a particular state that depends only on the initial inverse and forward models and on the supplied sequence of targets. Noise is not only present but necessary for the convergence of learning through the minimization of the difference between actual and predicted outcomes.

**Author summary:** Several studies have suggested that as we learn a new skill our brain forms representations, or “internal models”, of the skill and of the environment in which we operate. Theories of motor learning postulate that the brain builds forward models that predict the sensory consequences of motor commands, and inverse models that generate successful commands from planned movements. We test this hypothesis taking advantage of a special interface that generates a novel relation between the subject’s actions and the position of a cursor on a computer monitor, thus allowing subjects to control an external device by movements of their body. We recorded the motions of the body and of the cursor, and obtained estimates of both forward and inverse models. We followed how these estimates evolved in time as subjects practiced and acquired a new skill. We found that the description of learning as a simple deterministic process driven by the sequence of targets is sufficient to capture the observed convergence to a single solution of the inverse model among an infinite variety of alternative possibilities. This work is relevant to the study of fundamental learning mechanisms as well as to the design of intelligent interfaces for people with paralysis.

## Introduction

A distinct feature of the neuromotor system is the large number of muscles and degrees of freedom allowing it to attain a specific motor goal in a number of different ways [1]. This is both a resource and a computational challenge: while this motor redundancy provides the brain with a multitude of options, an enabling feature of motor dexterity, it also results in a family of ill-posed problems characterized by a lack of uniqueness in their solutions [2, 3]. Here, we consider the challenge posed by redundancy from the perspective of learning. How does the central nervous system learn to perform a novel task when multiple alternative solutions are available? This question acquires clinical relevance when a person suffering from loss of limb or some form of paralysis must reorganize the still available mobility to recover quality of life and independence through the operation of assistive devices – such as wheelchairs or robotic assistants – and dedicated human-machine interfaces.

In the last two decades, studies of motor learning [4-8] have established that the adaptation of limb movements to external perturbing forces takes place through the gradual formation of an internal representation, or “internal model” of these forces. To be predictable, the forces cannot be random disturbances, but must have a deterministic structure expressed in relation to the motion of the body and to the brain’s commands[6-9]. Donchin and Shadmehr [10] and others [11-13] have proposed to represent the development of such an internal model as the evolution of a dynamical system.

Internal models are of two types: forward and inverse. Forward models owe their name to their predictive representation of the process that transforms action commands into their sensory consequences. Inverse models reverse the direction of this process by deriving action commands from desired sensory outcomes. Earlier theoretical work by Jordan and Rumelhart [14] considered how the learning of actions can be viewed as the concurrent learning of forward and inverse models of actions. Here, we extend this approach to the learning of a novel map established by a body machine interface (BoMI) that translates movements of the upper body (shoulders and arm) into movements of an external object that users must guide to a set of target locations. This is an assistive tool for people that have lost the use of their hands after injury to the cervical spinal cord. We investigate how unimpaired subjects become skilled at controlling the external object via the BoMI. Our findings reveal that learning proceeds through the concurrent evolution of coupled forward and inverse models of the body-to-object mapping established by the BoMI. The validity of this description is tested by comparing the predicted evolution of motor performance with the learning performance observed in a group of human subjects. We compared the forward and inverse models derived from simulated learning dynamics with forward and inverse models estimated from motion data at different stages of learning.

## Results

### A model of learning while practicing control via a body-machine interface

We investigated how users of a body machine interface learn to reorganize or “remap” their body motions as they practice controlling an external object through the BoMI. The controlled object could be a wheelchair, a robotic assistant, or a drone [15-17]. Here we focus on the control of a computer cursor whose two-dimensional coordinates determine its location on a computer screen. Effectiveness in cursor control is the first and most common benchmark for brain-based interfaces [18-20], as the ability to control two-dimensional position is readily applied to a variety of tasks (e.g., an action performed via a joystick, entering computer text, etc.). We consider interfaces in which a linear mapping associates the body motion signals to the coordinates of the external object. Importantly, there is an imbalance between the dimensionality of the task space and that of the body signals, the latter being larger. Thus, any position of the controlled object corresponds to many (potentially infinite) different body configuration signals. The BoMI matrix *H* establishes a linear map between these two spaces; *H* has as many rows *K* as signals are needed to control the external object, and as many columns *S* as there are body signals. Not being square, the matrix *H* does not have a unique inverse. But there exist infinite “right inverses” that combined with *H* yield the *K*×*K* identity matrix in the task space of external control signals. Each such right inverse transforms a desired position of the controlled object in one particular set of values for the body signals. We consider users to be competent when they are able to move successfully their body in response to a presented target for the controlled object. Mathematically, we consider this as finding one right inverse *G* of the mapping *H*, out of a multitude of possible and equally valid choices. Current theories and experimental observations [10] suggest that learning is a dynamical process in which the learners modify their behavior based on the errors observed at each iteration of a task. In the kinematically redundant conditions considered here, learning is problematic because a given low-dimensional task error signal has multiple representations in the high-dimensional body space. Therefore, we considered an error surface defined by the squared task error in the space of the elements of the target-to-body map *G* adopted by a learner, where *G* is the learner’s inverse model of the body-to-cursor mapping *H* established by the BoMI. We implemented our learning model based on the hypothesis that the learners update the map *G* by moving along this error surface, following the line of steepest descent determined by the gradient of the squared error with respect to the elements of *G*. This error gradient depends on several variables; some can be directly observed by the learner, such as the error made in attempting to reach a given target position of the external device. However, the error gradient also depends upon the elements of the interface map *H*, which the learner cannot be assumed to know. Therefore, gradient descent learning of the inverse model requires a concurrent learning of the forward model. The latter requires a different error surface, since the forward map relates body motions to the consequent motion of the controlled object. Forward model learning does not require a target position for the controlled object, as the relevant error in this case is the difference between predicted and observed position of the controlled object. The squared prediction error defines an error surface in the space of the elements of the estimated forward map *Ĥ*.

Learning is thus described through two first order dynamical processes determined by two state equations. A forward learning process:

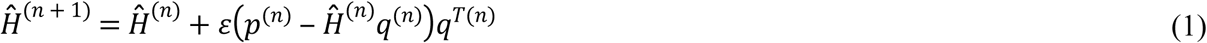

and an inverse learning process

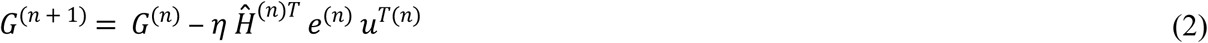

For details on the derivation of these equations, see Methods. The forward and inverse models are effectively the states of the respective processes, and the *n*-th iteration of the learning process results in state variables *Ĥ*^(*n*)^ and *G*^(*n*)^.

Equation (1) updates the subject’s estimate *Ĥ* of the forward model *H* that transforms *S*-dimensional body configurations *q* into *K*-dimensional positions *p* of the controlled object. The term in parentheses is the prediction error between actual and predicted positions of the controlled object. The concurrent process described by Equation (2) is the learning of the inverse model *G* that the subjects use to map the target object position *u*^(*n*)^ onto body signals. The reaching error *e*^(*n*)^ = (*p*^(*n*)^ − *u*^(*n*)^) that guides this process is the difference between actual and desired positions of the controlled object. Two possibly different learning rates, *ε* and *η*, provide inverse time constants for the respective dynamical processes.

Since we focus on the case in which forward and inverse learning are carried out concurrently, naïve users are immediately presented with the reaching task, and as they practice they observe both the reaching error and the prediction error. Equations (1) and (2) are coupled through *Ĥ*^(*n*)^ and through a third equation that describes the body signals currently adopted to reach the target position:

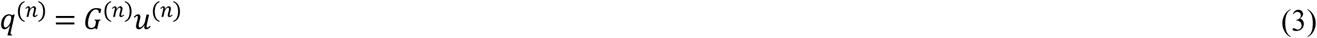

This apparently innocuous interaction has potentially harmful effects on the convergence of the coupled dynamics, as the second term in the gradient contribution to Equation (1) includes a quadratic factor in *q*^(*n*)^ and thus in *G*^(*n*)^. This contribution may result in local minima, a problem avoided by adding noise to Equation (3).

We validated our approach with six healthy subjects that learned to control the two-dimensional movement of a cursor on a monitor using eight signals from their upper body motions (shoulders and upper arms on both sides). In these experiments *S*=8 and *K*=2, and the state space of the combined forward-inverse learning was 2×16=32-dimensional.

### Dynamics of learning in human subjects

We monitored the learning process through two scalar metrics: RE, the L2 norm of the reaching error (the difference between actual and target location of the cursor at the end of the reaching movement), and IME, the spectral norm of the inverse model error (the difference between the identity matrix and the product between the interface map *H* and the current estimate*G*^(*n*)^ of the inverse model). The spectral norm of a matrix, indicated here by ‖ · ‖ to emphasize its analogy with the L2 norm of a vector, is the maximum singular value of the matrix. We estimated *G*^(*n*)^ from target and body signal data by least squares fit on Equation (3). The elements of *G*^(*n*)^ were estimated using data from 12 trials: trial *n* and its 11 preceding trials. Overlapping moving windows that included 12 trials were shifted by one trial at each iteration.

In the experiment, each subject practiced with a personalized body-to-cursor map derived from the statistics of its own freely produced upper body motions (see calibration procedure in Methods). With increasing number of practice trials, their RE decreased to values closer to the target radius (1 cm, Fig 1a). The learning process took over one hundred steps (172 ± 32 trials, mean ± SEM) before reaching asymptotic performance (Fig 1), identified as the time when the norm of the reaching error was smaller than the radius of the target. Similarly, the matrix*G*^(*n*)^ converged to a generalized inverse of the body-to-cursor map (Fig 1b). Although the subjects explored a number of different body configurations while learning how to control the cursor, in the end they found a stable movement pattern and built a representation of the inverse model *G*. Note the asymptotically small variations in the acquired *G*, with ‖Δ*G*‖ about 10% of ‖*G*‖ (Fig 2). For each subject, both RE and IME errors decreased with time following a trend captured by an exponential curve (Equation (21)). The corresponding learning rates, given by the inverse of the time constant of the exponential fits, are shown in Table 1 for each subject. Note the great similarity of these two rates for any given subject.

**Table 1.** Exponential rate used to best approximate the decay of RE (***λ***^***RE***^) and IME (***λ***^***IME***^) with ***n***, respectively. The R^2^ values quantify the goodness-of-fit of the exponential model for each subject to the corresponding experimental data.

| Subject | RE |  | IME |  |
| --- | --- | --- | --- | --- |
| | $\lambda^{RE}$ | $R_{RE}^2$ | $\lambda^{IME}$ | $R_{IME}^2$ |
| S 1 | 0.036±0.003 | 0.87 | 0.036±0.004 | 0.78 |
| S 2 | 0.010±0.001 | 0.85 | 0.006±0.001 | 0.86 |
| S 3 | 0.015±0.003 | 0.73 | 0.014±0.003 | 0.75 |
| S 4 | 0.016±0.003 | 0.80 | 0.022±0.004 | 0.80 |
| S 5 | 0.021±0.001 | 0.87 | 0.020±0.002 | 0.86 |
| S 6 | 0.029±0.002 | 0.93 | 0.029±0.002 | 0.90 |

**Fig 1.**
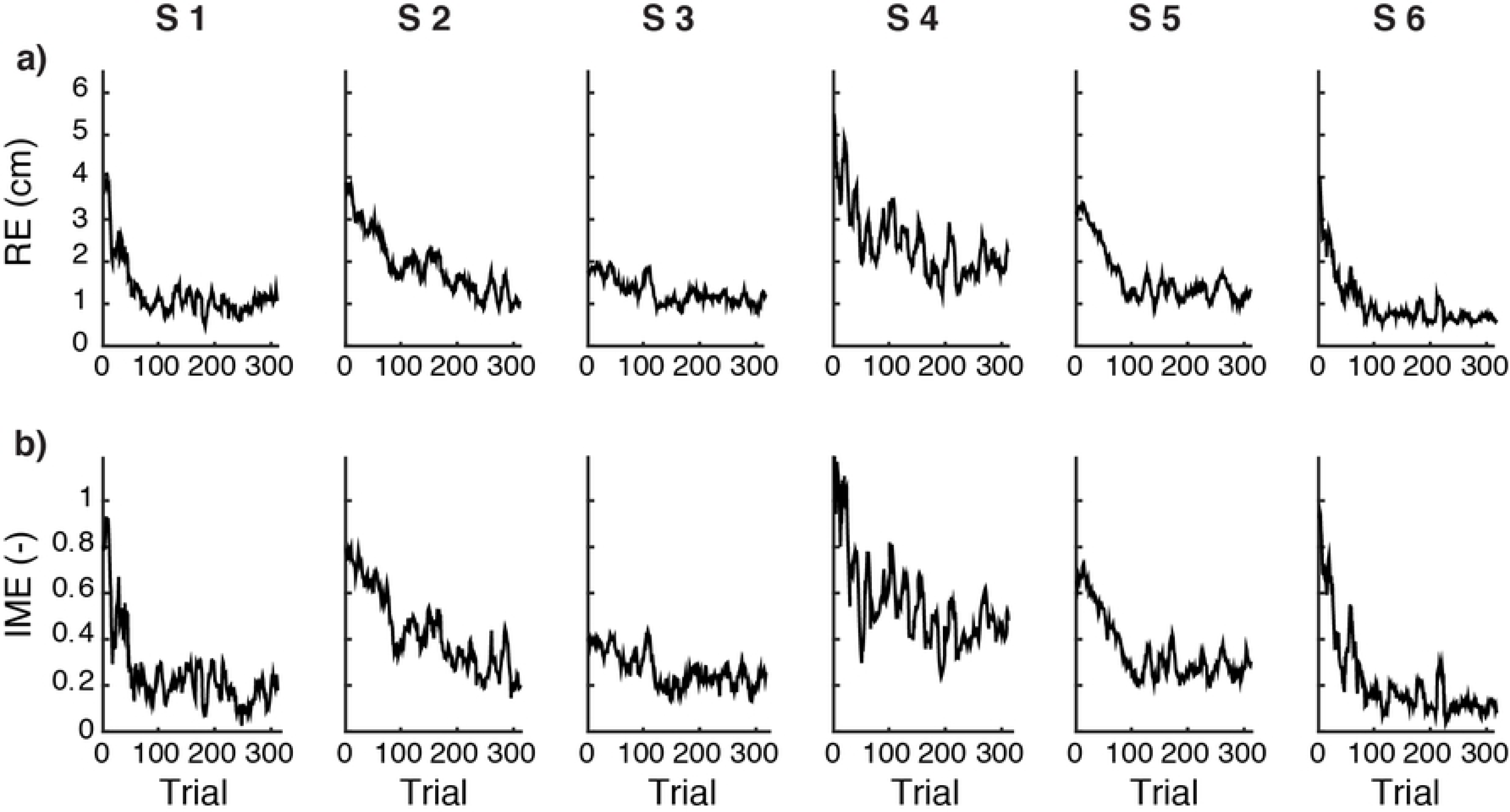
Subjects learn to use the body-machine interface. Data for the six subjects enrolled in the study (S1-S6). (**a**)Temporal evolution of the norm RE of the reaching error calculated over a moving window that includes the current and the 11 preceding trials. (**b**) Temporal evolution of the norm IME of the inverse model error. This includes the calculation of the inverse model ***G***^(***n***)^; the latter was obtained by least squares fit on Equation (3) from target and body signal data for trial ***n*** and the 11 trials preceding it.

**Fig 2.**
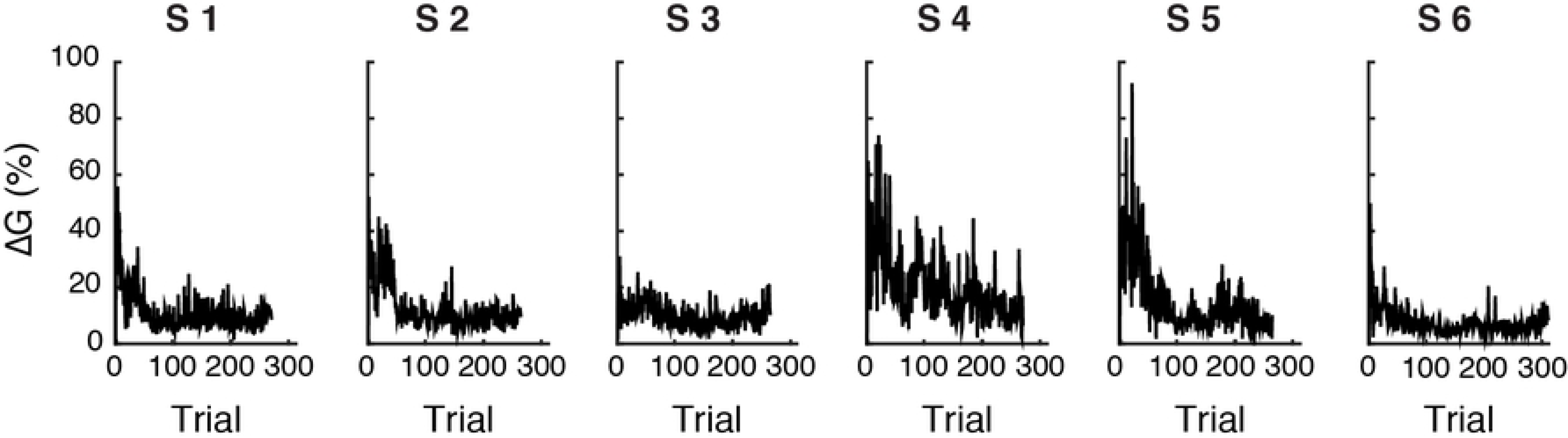
Temporal evolution of the changes in the inverse map *G*^(*n*)^. Changes in the inverse map as a function of trial number ***n***, quantified by ‖**Δ*G***^(***n***)^‖ = ‖***G***^(***n***)^ − ***G***^(***n*** − **1**)^‖ /‖***G***^(***n***)^‖(see Equation (18) in Methods), for the six subjects enrolled in the study (S1-S6).

### Learning dynamics: model vs. human subjects

To build a model for each subject, we used the subject-specific map *H* and the subject-specific sequence of targets used for training. The learning rate *η* for the inverse model (Equation (2)) was taken to be equal to the subject-specific rates *λ*_*RE*_ reported in Table 1 (see Methods). The learning rate *ε* for the estimation of the forward model (Equation (1)), and the amplitude *σ* of the noise added to the inference of body motions (Equation (3)) followed from an optimization procedure (see Methods). Table 2 reports the values of these three parameters for each subject. We let the model evolve until the norm of the reaching error was smaller than 1 cm, as the subject’s performance reached a plateau when the cursor reached the target.

**Table 2.** Subject-specific model parameters. The learning rates ***η*** and ***ε*** correspond to the inverse and the forward model, respectively; ***σ*** is the amplitude of the Gaussian noise added to the inference of target-specific body motions.

| Subject | $\sigma$ | $\varepsilon$ | $\eta$ |
| --- | --- | --- | --- |
| S 1 | 0.7335 | 0.1774 | 0.036 |
| S 2 | 0.6587 | 0.2463 | 0.010 |
| S 3 | 0.7395 | 0.1924 | 0.015 |
| S 4 | 0.7241 | 0.2178 | 0.016 |
| S 5 | 0.8278 | 0.1540 | 0.021 |
| S 6 | 0.7794 | 0.1937 | 0.029 |

We then tested how well our model captured the learning dynamics of each subject. As shown in Fig 3, these subject specific models were able to predict quite well the individual learning curves. Both the RE and the IME estimated from the model follow the time evolution extracted from the real experimental data. Correlation coefficients (Table 3) quantify the similarity between the simulated and actual temporal evolution of RE and IME during learning.

**Table 3.**
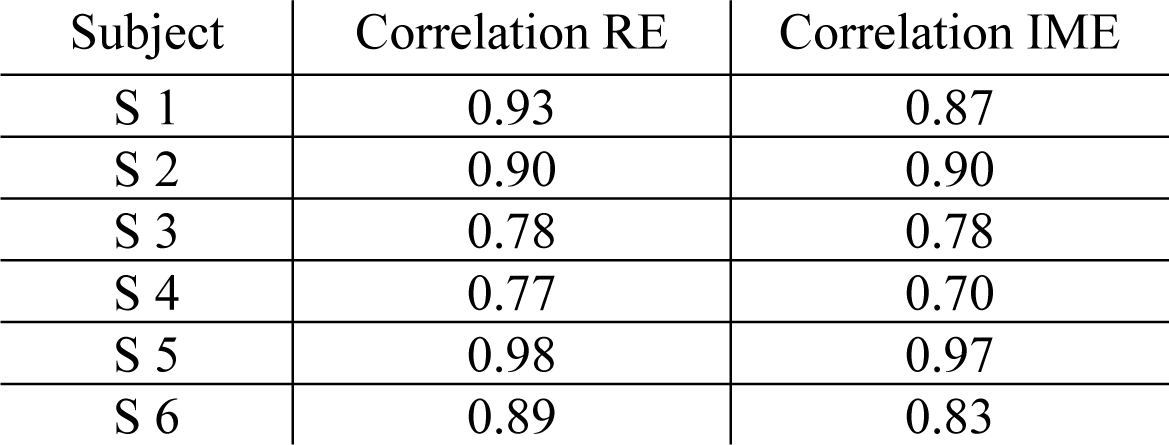
Correlation coefficients (R^2^) between the temporal evolutions of the norm RE of the reaching error and the norm IME of the inverse model error recorded during the experiment, and the temporal evolution of these two scalar error metrics as predicted by the model.

| Subject | Correlation RE | Correlation IME |
| --- | --- | --- |
| S 1 | 0.93 | 0.87 |
| S 2 | 0.90 | 0.90 |
| S 3 | 0.78 | 0.78 |
| S 4 | 0.77 | 0.70 |
| S 5 | 0.98 | 0.97 |
| S 6 | 0.89 | 0.83 |

**Fig 3.**
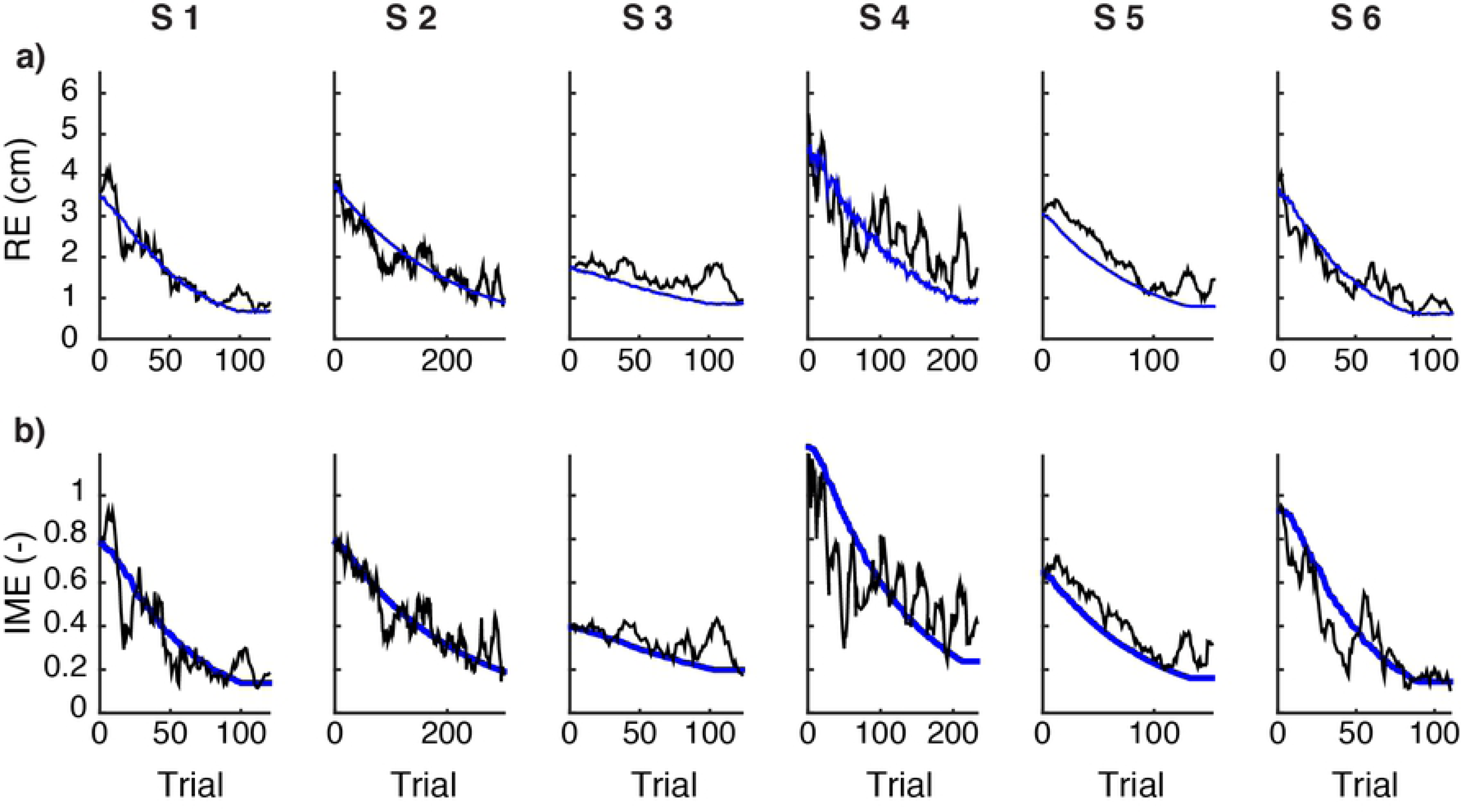
Modeling human learning. (a) Temporal evolution of the norm RE of the reaching error as a function of trial number ***n***, calculated from the data of six subjects (black) and from the respective models (blue). (b) Temporal evolution of the norm IME of the inverse model error as a function of trial number ***n***, estimated from the experimental data (black) and from the model simulation (blue). Both metrics are calculated over a moving window that encompasses trial ***n*** and its preceding 11 trials.

The model also allows us to compute a current estimate of the forward map *Ĥ* as acquired by the subjects while practicing. We quantified the similarity between the estimated *Ĥ*^(*n*)^ and the BoMI map *H* at each iteration of the learning dynamics by using the forward model error (FME), defined as the spectral norm of the difference between *H* and *Ĥ*^(*n*)^, normalized by the spectral norm of *H* (see Equation (19) in Methods). This error in the estimate of the map that transforms body movements into cursor movement was responsible for the cursor prediction error, the difference between the actual and the expected position of the cursor. We monitored the norm PE of the prediction error and the FME as a function of trial number *n* (Fig 4). The estimate *Ĥ*^(*n*)^ converged toward the actual forward map *H*, resulting in near zero asymptotic values for both PE and FME (Fig 4).

**Fig 4.**
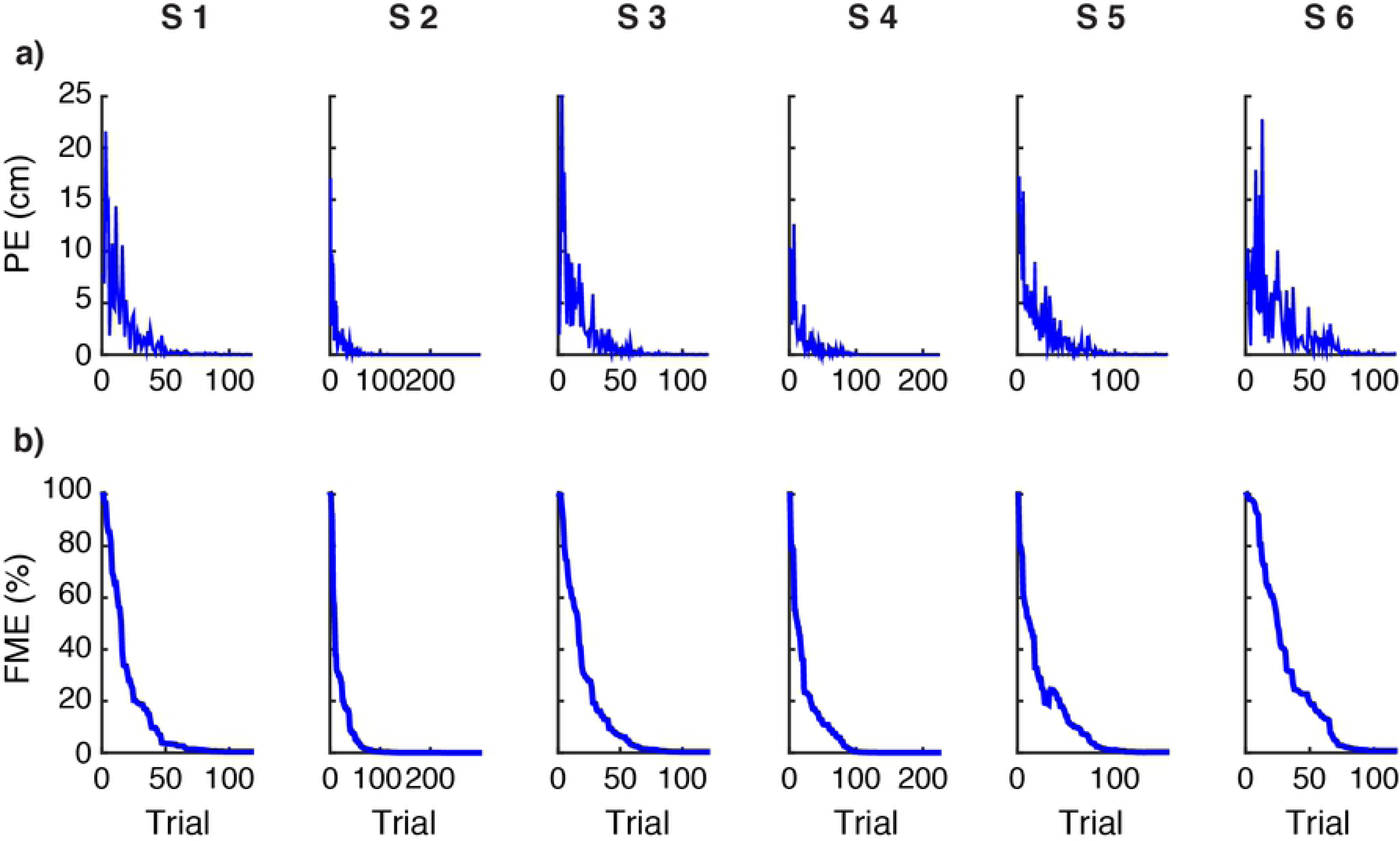
Subjects learn the forward model. (**a**) Temporal evolution of the norm PE of the prediction error as a function of trial number ***n***. (**b)** Temporal evolution of the estimate of the forward map, quantified by FME = ‖ ***Ĥ***^(***n***)^ − ***H***‖/‖***H***‖, as a function of trial number ***n***. Data are shown for each subject-specific model. Only the first 100 iterations of the dynamics are presented, since the asymptotic regime has by then been reached for both metrics.

The dynamics of the learning model captured the errors in the low-dimensional task space of the controlled cursor as well as the history of body signals generated by the subjects in response to the successive targets (Fig 5 and Table 4); the sole exception was Subject 4, whose accuracy in reaching the target position was smaller and characterized by a higher level of variability (Fig 3a). The body and cursor signals recorded during the experiment and predicted by the model were not very similar at the beginning of training, but they quickly converged and tended to overlap by the time RE and IME reached their asymptotic condition (Fig 5, Table 4). These results support the conclusion that a model of learning based on simple gradient descent over quadratic error surfaces captures the formation of forward and inverse representations of the map established by the body-machine interface.

**Table 4.** Correlation coefficients (R^2^) between the temporal evolution of the body signals {q_1_, …, q_8_} and the cursor coordinates {p_1_,p_2_} recorded during the experiment, and the temporal evolution of these quantities as predicted by the model. The comparison was performed during the last 100 trials.

| Subject | Body space | Cursor space |
| --- | --- | --- |
| S 1 | 0.90 | 0.96 |
| S 2 | 0.90 | 0.97 |
| S 3 | 0.93 | 0.98 |
| S 4 | 0.69 | 0.93 |
| S 5 | 0.91 | 0.98 |
| S 6 | 0.93 | 0.99 |

**Fig 5.**
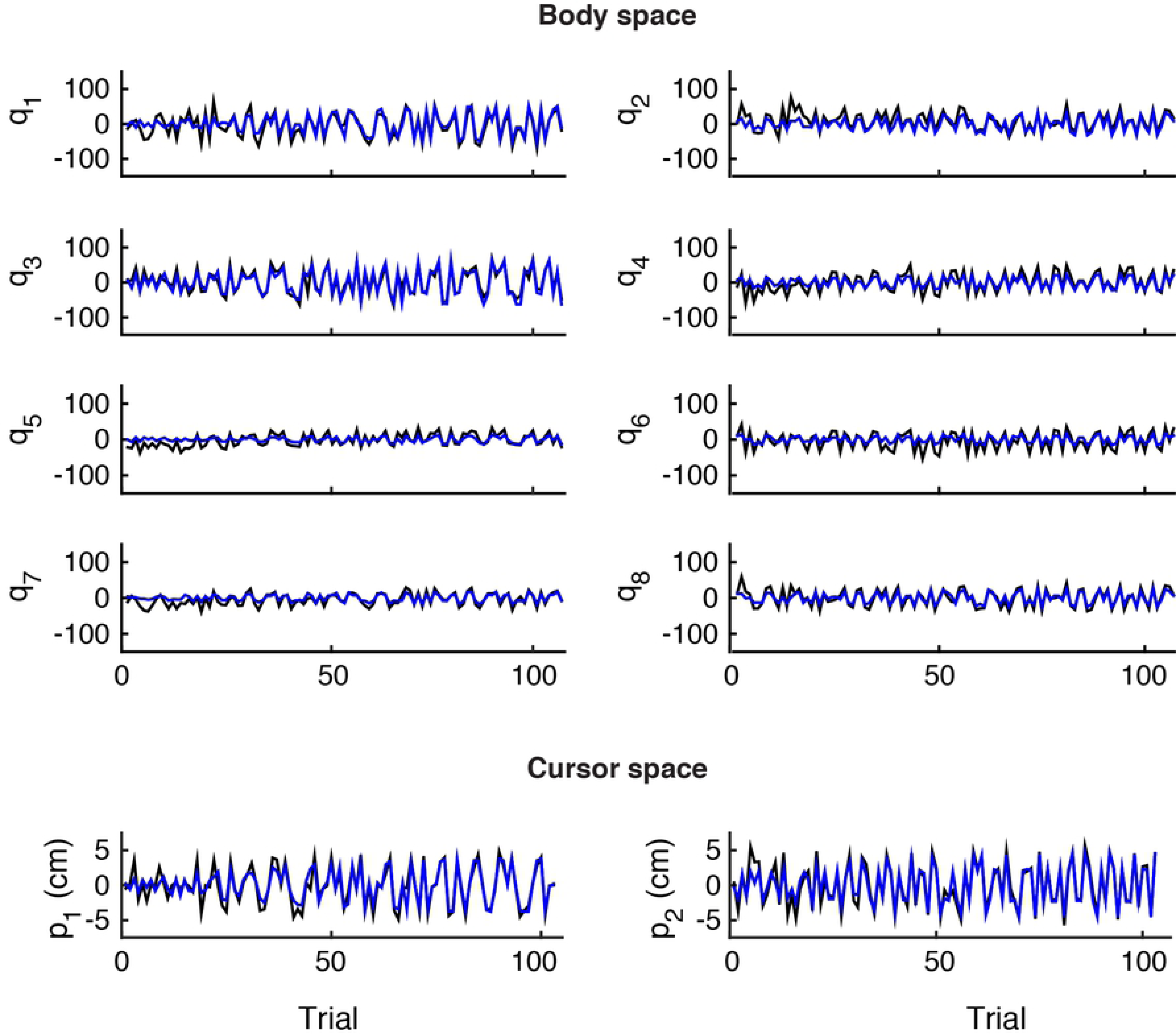
Reconstruction of body and cursor signals. Comparison between real (black) and simulated (blue) data for subject 6. The top panel presents the values of the eight body signals {q_1,_…,q_8_}, i.e., the (x,y) coordinates of four markers (shoulders and upper arms on both sides) in the image plane of the associated cameras; the bottom panel shows the (x, y) coordinates of the cursor {p_1_, p_2_}in the reference frame of the computer monitor.

## Discussion

We investigated the learning process that occurs when subjects reorganize or “remap” their body motions as they learn to perform a task that involves a novel relation between body motions and their observable consequences. For a patient who suffers from severe paralysis and is obliged to reorganize the available mobility to operate an assistive device such as a powered wheelchair through a human-machine interface, the ability to engage in such remapping becomes a necessity of life. Here we investigated the process of learning to perform reaching movements via a body-machine interface; we used a group of unimpaired subjects under the preliminary assumption that similar learning mechanisms are present in people suffering from injuries to the spinal cord. We considered a body-machine interface that harnesses signals generated by body motions in order to control an external object, in this case the position of a cursor on a computer monitor. The interface establishes a many-to-one body-to-object map that the user must learn to master starting from a naïve state. The map is in fact not intuitive, as there is no obvious correspondence between the motions of the body and the motion of the controlled object.

In our experiment, unimpaired subjects practiced controlling a computer cursor through a BoMI whose linear body-to-object map was customized to each subject through a calibration procedure that fitted the statistics of the subject’s body motions. Through practice with a fixed BoMI, all subjects demonstrated exponential convergence toward an inverse of the BoMI mapping, with subject-specific learning rates. The parameters of this inverse model define a state space in which learning is modeled as a first order dynamical process that evolves based on the specific sequence of target positions for the external object, and of two types of error: (i) the prediction error that is the difference between the actual position of the object and the position predicted by the subjects based on their internal representation of the interface map, and (ii) the task error or reaching error that is the difference between the position reached by the object and the actual target position. While the prediction error depends on the current estimate of the forward model, the reaching error depends on the current state of the inverse model, which determines the chosen body motion.

We studied the evolution of the learning system using different interface maps *H*, learning rates *η*, and target sequences *U* for each subject. The empirical observations of human adaptation to the BoMI were compared to predictions from a subject-specific learning model that used initial conditions inferred from each subject’s initial performance, the same *H* and *U* as used by the subject, and a learning rate *η* obtained by temporal regression of the experimental data. We demonstrated that a model based on first-order dynamics was sufficient to capture the evolution of learning as described both through the observed errors and through the accuracy of the estimated internal models. There was however a notable difference between the models’ and the subjects’ learning: the decay of the norm of the reaching error computed from real data was not as smooth as the decay predicted by the model. This might be due to the main simplifications adopted in our model, namely, i) the learning dynamics (Equations (1) and (2)) were assumed to be deterministic, without any noise contributing to the state and output equations, and ii) the learning of the inverse model (Equation (2)) was assumed to be linear.

Although the interface forward map is linear (Methods, Equation (4)), it is a many-to-one map admitting a multitude of inverses. This “redundancy” opens the possibility of successful linear and nonlinear inverse maps. Redundancy also leads to an important consideration about gradient descent learning. The reaching error surface in the space of the inverse model elements does not have a unique minimum but a continuously connected set of minima corresponding to the null space of the forward map. In the metaphor of a skier descending from a mountain following the gradient, this space of equivalent inverse models corresponds to a flat elongated valley at the bottom of the mountain. Anywhere along the valley is a valid end to the ride, as it corresponds to a valid inverse model. The inverse model on which the steepest descent ends depends on the initial conditions, as predicted by the dynamical model (Fig 3b – evolution of the norm of the inverse model error).

The analysis of learning dynamics of the users of a body-machine interface is essential for the effective development of coadaptive approaches, where the interface parameters are themselves updated based on the user’s state of learning [21-23]. Coadaptation requires a seamless integration between machine learning and human learning. Both learning processes are dynamic, evolving as functions of their own internal state and of inputs that reflect the state of their counterpart. A mismatch between the timing of the interface updates relative to the subject’s learning dynamics would likely lead to hindering human learning (if the interface update rate is too fast) or to ineffectiveness (if the interface update rate is too slow).

In our current understanding, motor learning is not only a way to acquire or improve a skill, but is perhaps most importantly a biological mechanism to gain knowledge about the physical properties of the environment [9]. Through the practice of movements, the brain learns to separate unpredictable from predictable features of the world in which the body is immersed; through the formation of representations or “internal models” of the predictable features, the brain acquires the ability to anticipate the sensory consequences of its commands. The human operator of a BoMI must develop an inverse model of the forward map to transform a desired goal into an action of the body. Further evidence from the literature indicates that adaptive, error-driven, internal model formation is a general feature of motor learning, observed during arm reaching and drawing, and during pointing with the legs and walking [24].

Theoretical studies of motor learning have focused on control policies and internal models to understand how the brain generates action commands [25]. Control policies allow the brain to select goals and plan actions, while internal models generate motor commands that are appropriate for those plans in the context established by sensory feedback. For example, when the goal is to reach a target, the brain must first evaluate the current position of the limb with respect to the target and use a control policy to plan a movement of the hand [3, 26]. An internal model of limb’s dynamics, called an inverse model, then converts that plan into motor commands [27-29], while a forward model of the same limb’s dynamics predicts the sensory consequences of these motor commands [14, 30, 31]. The comparison of this prediction with actual sensory feedback [32] allows re-estimating the current hand position with respect to the target and updating the motor plan [33] by issuing an error-dependent motor command aimed at correcting the ongoing movement. Forward and inverse internal models thus play a fundamental role in movement planning and execution.

Recent studies have considered the formation of these internal models as dynamical process [10, 34, 35]. For example, to account for findings observed when reaching arm movements are perturbed by external force fields, Donchin and co-workers [10] argued that the forces generated by the subjects to compensate for the external field are the output of an internal model of the field, developed through experience. Their theory, successful at predicting the time history of adaptation, was based on two key assumptions that we also adopted here, namely i) that the movement outcomes and the ensuing errors result from a deterministic process, and ii) that the internal model is the state of learning.

The transformation from the movements of the BoMI user to the movements of the controlled objects establishes a new geometrical relation between body motions and their consequences. The BoMI thus essentially creates a novel geometry that the user must learn to operate. Here, we have implemented a linear transformation from body signals to a cursor, which allowed us to work under the assumption that the users would develop a linear inverse model of this map. However, linearity of the inverse map is not a necessary consequence of operating through a linear forward map, because a linear forward map that is not bijective may also admit nonlinear inverses. Therefore, our approach will not result in the most general solution to the problem of finding an inverse map. Nevertheless, our analysis demonstrated that the linear inverse model derived by coupled gradient descent on both the prediction error and the reaching error is capable of reproducing with high fidelity the entire history of a subject’s responses to a sequence of targets (Fig 5). While we are unable to exclude more complex processes that could lead to an equally effective nonlinear inverse model of the linear BoMI map, the linearity assumption not only leads to results that agree with the experimental data but also fulfills Occam’s razor criterion for simplicity.

We conclude with some comments on the clinical relevance of this study. Damage to the spinal cord, stroke, and other neurological disorders often cause long-lasting and devastating loss of motion and coordination, as well as weakness and altered reflexes. In most cases, some residual motor and sensory capacities remain available to the disabled survivor, and can be harnessed to provide control signals to assistive devices such as robotic systems, computers, and wheelchairs. A first challenge for the disabled [36] is to learn how to interact with the assistive devices and how these respond to the user’s actions.

A broad spectrum of sensors, such inertial measurement units (IMUs) placed on the head [37] or electroencephalography (EEG) systems [38], are available for detecting and decoding movement intentions. A body machine interface (BoMI) captures residual body motions by optical [17, 39, 40], accelerometric [15, 41], or electromyographic sensors [42], and maps the sensor signals onto commands for external devices such as powered wheelchairs [15] or drones [17], or onto computer inputs. At the other end of the spectrum, brain-machine interfaces decode motor intention from neural activity recorded in motor or premotor cortical areas [19, 20, 43]. Both brain- and body-machine interfaces take advantage of the vast number of neural signals and degrees of freedom of the human body [1, 44, 45], and of the natural ability of the motor system to reorganize the control of movement [4, 9, 46, 47]. Typically these interfaces establish a map - most often linear - from the space of neural or motion signals to the lower dimensional space of control signals for the external device [41, 48]. The user’s ability to operate the interface is expected to change over time; either a positive change associated with the acquisition of greater control skills, or a negative change due to the worsening of the user’s medical conditions. In either case, the interface map also needs to change, to coadapt with its user. This coadaptation is a critical challenge in the development of both brain- and body-machine interfaces [21, 23]; harmonizing the interface update with the processes that guide the improvement or decay of the user’s skill is of obvious importance. Understanding the dynamics of human learning through the interaction with the interface carries the promise of creating truly intelligent systems capable of compensating for the changing abilities of their users [19, 22].

## Methods

### Computational model

Inverse kinematics is a well-known and well-explored computational problem in robotics [49, 50] and human motor control [27, 51]; it refers to finding the configuration of joint angles that results in a desired position of an end effector or of the hand in the operational space [52]. Inverse kinematics problems become ill-posed [53] when there are multiple valid solutions as a consequence of the many-to-one nature of the forward kinematic map. This is the situation considered here, in which the kinematics that the subjects are controlling may be partitioned in a sequence of two maps. In a first map, the subjects control the motions of their bodies by acting on a multitude of muscles and joints. In a second map, the signals triggered by these body motions determine the lower dimensional state of an external object such as a wheelchair [15], a cursor on a computer screen [16], or a drone [17]. Here we make the critical but reasonable assumption that the subjects have already acquired in a stable form the expertise needed to control the motion of their body, or at least portions of it that were unaffected by injury or disease. Therefore, they only need to acquire the second component. This is the component we focus on, limited here to a body-machine interface whose linear map *H* transforms, at any given trial *n*, an *S*-dimensional vector of body signals *q* into a *K*-dimensional control vector *p* as follows:

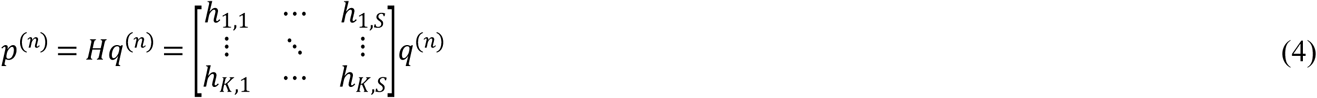

Here, *H* is a *K*×*S* matrix. Since *K*<*S*, this interface map is many-to-one and there is a “null space” of inputs for each value of its output, encompassing all different patterns of body signals that result in the same control signal. This is an important characteristic of the map; earlier work has shown that subjects learn through practice to separate the null space from its orthogonal “potent space” complement [48].

#### Learning dynamics as first order state-based model

In a learning experiment where the goal is to reach targets in the control space, the superscript *n* labels the trials or successive repetitions of a single action; for instance, each trial is a reaching movement in a sequence of such movements. At the end of a trial, the learner observes an error *e*^(*n*)^. This error drives the updating of the internal model, which we assume to be a liner map *G*^(*n*)^ transforming a goal *u*^(*n*)^ into its corresponding body vector (previously Equation (3)):

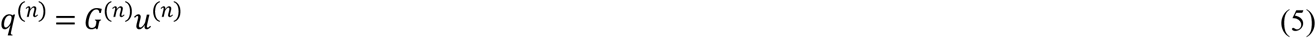

Since the forward map *H* is linear, the linearity of *G* is a sufficient but not necessary condition. More complex nonlinear structures of the inverse model would in principle be admissible. Here, we assume the simplest general form for a linear inverse model of the forward BoMI map; this assumption makes the investigation of learning dynamics tractable.

For the *n*-th reaching trial, *u*^(*n*)^ is the position of the target, and the reaching error is the *K*-dimensional vector from the target position to the actual position of the controlled object at the end of the trial:

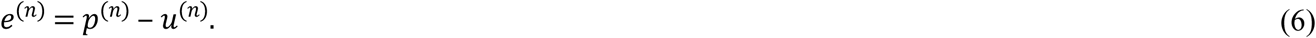

As learning reaches a steady state, participants are expected to have eliminated this error. This implies that the internal model becomes a right-inverse model of the interface map: since

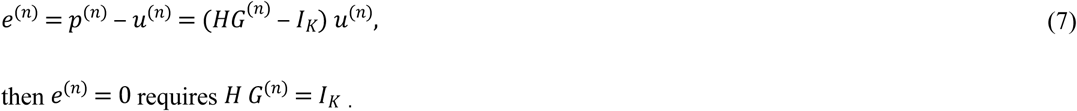

Learning is represented as a dynamical process whose state, the internal inverse model *G*^(*n*)^, changes after the observation of each reaching error. The targets presented to the learner constitute the external input to this process. To ensure that the change in state leads to a reduction of the error, the learning process drives the state along the gradient of the quadratic error surface in the state space defined by the components of *G*. The gradient of the squared reaching error with respect to the components of the inverse model is

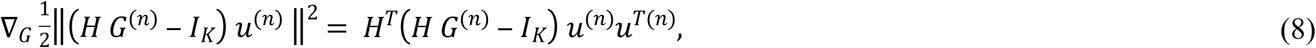

which leads to the update equation

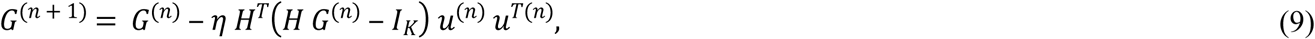

or, equivalently

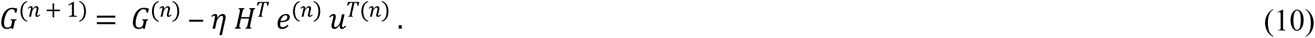

Here, *η* is a learning rate parameter that we model as a scalar, although in principle there could be a different rate for learning every element of the forward and inverse models. We found that only two learning rates, *ε* for the forward model and *η* for the inverse model, sufficed to account for the observed learning behavior.

If the interface map *H* is known, the update Equation (10) provides an estimate of the right inverse of *H* solely on the basis of control space data, without performing an explicit matrix inversion. Given the targets *u*^(*n*)^, the variables of interest are the observed reaching errors *e*^(*n*)^ and the estimated inverse model or “state of learning” *G*^(*n*)^. As *e*^(*n*)^→0, *G*^(*n*)^ becomes stationary.

The gradient of the error involves the actual value of the interface map *H*. It is not plausible to assume that our subjects had any initial notion of the interface map, let alone an exact representation. In a realistic model of learning, the value of *H* must be replaced with an evolving estimate *Ĥ*^(*n*)^. In this scenario, the current state of learning is represented in a higher dimensional space that includes the components of both *G*^(*n*)^ and *Ĥ*^(*n*)^. In the case of *S*=8 body signals controlling the location of an object in *K*=2 dimensions, the state space for learning is 2×8×2=32-dimensional. We follow a concept introduced by Jordan and Rumelhart [14], and represent learning as the parallel development a of forward-inverse model.. The forward model leads to a prediction of the controlled device position given the current set of body signals, whereas the inverse model generates the body signals needed to achieve a given target position of the device.

In principle, the two learning processes could take place in two separate phases: a “flailing” phase where aimless body motions are produced and the resulting object motions are compared with their expected motion to obtain prediction errors that drive the estimation of the forward model. This would then be followed by a phase where the subjects reach for specific targets, and reaching errors drive the estimation of the inverse model (see S1 Fig). This mechanism has been suggested as a possible model of motor development in infants [54], who acquire a model of the dynamical properties of limbs by driving them with erratic neuromuscular activities. However, this is not the case in experiments where subjects are presented with reaching targets from the onset; in this scenario, no aimless “flailing” was observed. It is thus more plausible to model forward and inverse learning as concurrent processes.

The prediction error quantifies the difference between actual and predicted positions of the controlled object, without reference to a target. The gradient of the squared prediction error with respect to the components of the forward model *H* is

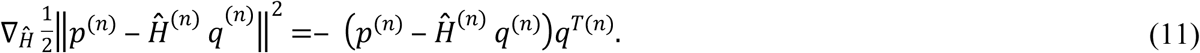

The update equations for the coupled learning process then are

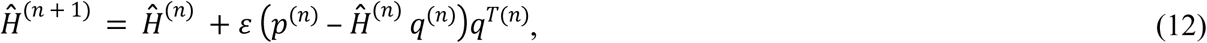

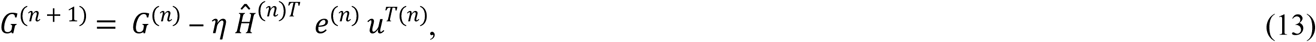

reported earlier as Equations (1) and (2). Note that Equation (12) contains a term that is quadratic in *q*; this implies a quadratic dependence on the elements of *G*^(*n*)^, since at each step *q*^(*n*)^ is derived by applying *G*^(*n*)^ to the corresponding target. The update equation for the estimate of the forward model is thus not linear, and the learning process is prone to get stuck in local minima. The simplest way to avoid this is to add a small amount of noise to the body signals derived from the inverse model,

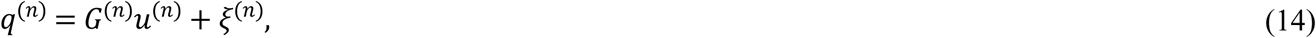

where the noise *ξ* has the same dimension *S* as *q*, and each component of *ξ* is independently drawn from a Gaussian distribution *𝒩*(0, *σ*^2^) at each trial.

Two important parameters of the combined learning model of Equations (12) and (13) are the learning rates for the forward (*ε*) and the inverse (*η*) models. In the case of eight body signals mapped into a two-dimensional control space, these learning rates apply to the evolution of 16 elements each; their most general form would be a 2×8 matrix for *ε* and an 8×2 matrix for *η*. Here, for simplicity and to avoid overfitting, we assume both learning rates to be scalar. A similar assumption is made for the noise amplitude *σ*, a somewhat less critical parameter whose sole purpose is to add sufficient noise to the learning algorithm so as to avoid getting trapped in local minima.

### Validation of the model with experimental data

To validate the outcomes of the model we recruited six unimpaired subjects (age range 21-40 years old, 3 males and 3 females) in the preliminary study. All of them signed an informed consent approved by Northwestern University Institutional Review Board.

The subjects practiced the execution of reaching movements via an interface (Fig 6) that mapped an eight-dimensional signal space associated with upper body motions to the two-dimensional task space of a computer cursor. An array of four video cameras (V100, Naturalpoint Inc., OR, USA) was used to track active infrared light sources attached to the subject’s upper-body garments (two for each side of the body, one on the shoulder and one on the upper arm, as shown in Fig 6). Each camera pointed at a single marker, providing two signals defining the coordinates of the marker in the camera’s frame. Collectively, the four sensors provided an eight-dimensional body vector *q* that was transformed into a command vector *p* for controlling the position of a cursor on a computer screen, *p* = *Hq*.

**Fig 6.**
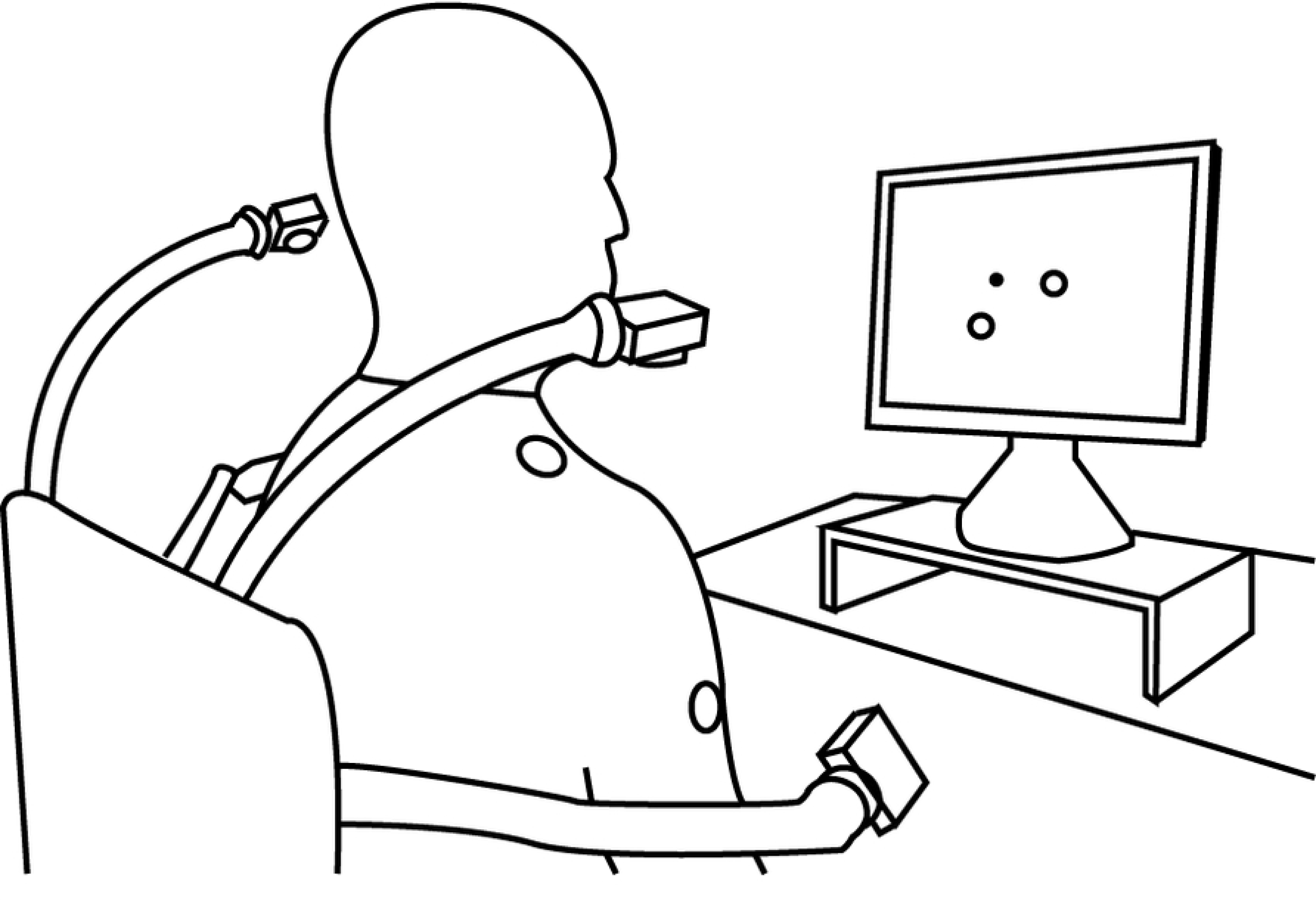
Experimental setup of the Body Machine Interface. The subject sits on a chair in front of the computer. Active optical markers were placed on shoulders and arms; their positions were recorded by infrared cameras.

In order to design the matrix *H* for each subject, we used a standard linear dimensionality reduction method, principal component analysis (PCA) [55] to set up the dimensionality reduction implicit in the *q* to *p* map. PCA is based on the decorrelation of the input signals through the diagonalization of their covariance matrix; dimensionality reduction is obtained by keeping only the eigenvectors corresponding to the *K*=2 largest eigenvalues. PCA provided us with a computationally straightforward method for identifying those directions that captured the largest extent (i.e. the largest variance) of body motions within the space of sensor signals for each subject.

The interface map *H* from the input space of body signals to the output task space was constructed in three steps:

1. Calibration – Subjects were asked to freely explore their range of shoulders and upper arms motion in all possible directions for about 60 seconds. This stage results in an aimless free dance, during which subjects visited a large portion of their available range of motion; their movements were limited only by the type and degree of impairment. The “calibration” data set *Q*_*CAL*_ was organized as an *S*×*M* matrix, where the number *S* of rows is the dimensionality of the space of sensor signals (here, *S*=8) and the number *M* of columns is the total number of samples (here *M*=5000, collected over about 66 seconds at 75 samples per second).
2. PCA – The principal components of the data set *Q*_*CAL*_ were extracted using PCA. The *S* eigenvectors of the covariance matrix of the sensor signals were ordered according to the magnitude of their corresponding eigenvalues, from largest to smallest. The eigenvalues represent the variance of the data along the eigenvector directions; those directions with high signal excursion correspond to larger eigenvalues. We considered those high-variance movement directions to be the user’s “best controlled” combinations, and used the *K* leading eigenvectors as rows to construct the *K*x*S* matrix *H* (Equation (4)). In our case the interface map *H* was a 2×8 matrix; the two first PCs accounted for 73±5% of the variance of the calibration dataset across subjects.
3. Training – After calibration, participants went through a two-hours training session. The training session consisted of 324 reaching movements from the center of the screen to targets located 5 cm away from the center in 6 directions 60° apart. Although the order of target presentation was randomized, a given target was not presented until the subject had reached all other targets. The subjects had no visual feedback about the cursor position for the first second following movement onset. The cursor then became visible, and the subjects could use visual feedback to correct for the reaching error if the cursor was not on the target.

### Data analysis

#### Estimation of the learning dynamics

To investigate the learning dynamics, we focused on the temporal evolution of two scalar variables in task space: the reaching error (RE) and the inverse model error (IME). These two variables were computed both from the data for healthy volunteers and from the synthetic data generated by simulating the proposed model for the learning process.

The reaching error RE was computed as the norm of the difference between the actual cursor position at the end of the reaching movement and the target position. For the experimental data, we considered as an estimate of the reaching error the distance between target and cursor at the end of the blind phase of the trial, when the subjects moved relying only on their inverse internal model in the absence of visual feedback of the cursor motion. As the cursor reappeared, the subjects performed a corrective movement bringing the cursor on the target. The inverse model error IME was computed as the norm of the difference between the identity matrix *I*_*K*_ and the product *H G*^(*n*)^ between the interface map and the estimate of the inverse map at the end of each trial.

The lower dimensionality of the output space for the interface map *H* causes the problem of finding the inverse map to be ill-posed; the surface defined by the squared reaching error in the state space spanned by the components of *G* does not exhibit a single minimum but a flat extended “valley” corresponding to all possible inverses of the interface map *H*. To circumvent this ambiguity and to monitor whether subjects converged towards a stable inverse transformation, we estimated the inverse model matrix *G*^(*n*)^ from the subjects’ performance.

A typical experimental data set consisted of temporal sequences of reaching movements. At each trial *n* we considered a movement set, a sequence of *r* trials that included the *n*-th trial and the (*r*-1) trials that preceded it. Here we used *r*=12, so that on average each movement set included two trials towards each of the six different targets. The body and target vectors for the *n*-th movement set were collected in the arrays

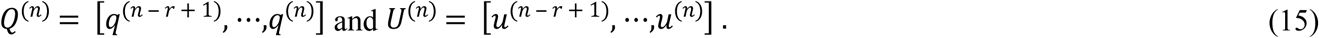

The matrix *G*^(*n*)^ then was obtained from a least-squares estimation based on Equation (3):

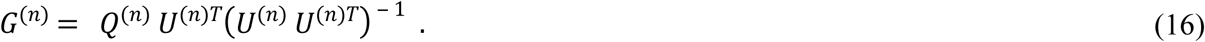

The history of reaching errors for the *r* trials in the movement set that ended with trial *n* was computed as

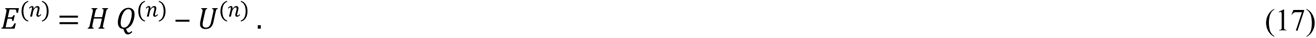

A scalar reaching error (RE) was then calculated by taking the spectral norm ‖*E*^(*n*)^‖ of the *K*x*r* matrix in Equation (17). Similarly, we calculated the inverse model error (IME) as the spectral norm ‖*H G*^(*n*)^ − *I*_*K*_ ‖. The IME is expected to approach 0 as learning converges and *G*^(*n*)^ approaches a right inverse of *H*. The convergence toward a stable representation of the inverse map was assessed by computing the percentage difference in norm among consecutive estimations of *G*^(*n*)^,

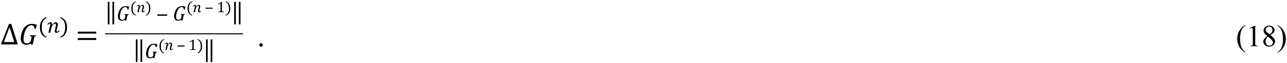

We defined two additional errors to quantify whether each subject was also forming a forward map *Ĥ* that converged to the interface map *H*. We defined the forward model error (FME) as

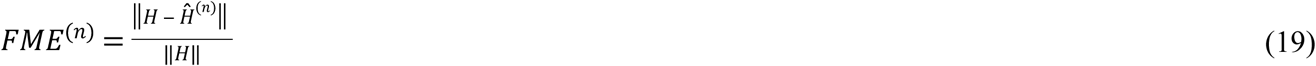

The current estimate *Ĥ*^(*n*)^ affects the prediction error, computed in task space as the difference between the actual position of the cursor and the one estimated with the current estimate of the forward model. The L2 norm of this difference defines

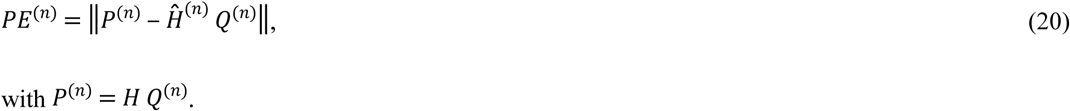

No moving window was used to compute these two errors because both PE and FME could be extracted from the simulated data at each trial.

#### Model parameters

For each subject recruited for the study, we constructed a model that used the same interface map *H* as used by the subject; the model was then exposed to the same target sequence. To set the individual learning rate *η* for the learning of the inverse map in Equations (2) and (13), we fitted the experimentally observed decay of the norm RE of the reaching errors for each subject to an exponential of the form

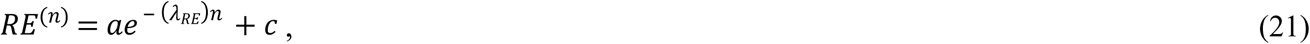

to obtain a value of *λ*_*RE*_ for each subject. We then set *η* = *λ*_*RE*_ in the corresponding subject-specific model.

To set values for the parameters *ε* and *σ* we adopted a minimum search approach to minimizing a cost function based on the forward model error (FME), as those two parameters mostly influence the evolution of the estimation *Ĥ* of the forward model. The cost function *C* was defined as

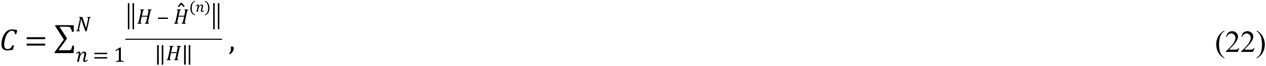

where *N* is the total number of trials.

#### Comparison between simulated and real data

To estimate the similarity between the real and model data we used the correlation coefficient R^2^. By definition the R^2^ evaluates similarity between the shapes of the compared curves and also provides additional information regarding the amplitude. This metric was applied to the norm RE of the reaching error, the norm IME of the inverse model error, the body coordinates *q*, and the cursor coordinates *p*. For each of these quantities, assume that the measured values are {*y*^(*n*)^}, 1 ≤ *n* ≤ *N*, where *N* is the total number of trials. The model gives a prediction or estimation {*ŷ*^(*n*)^}, 1 ≤ *n* ≤ *N* for each of these values, with residuals *e*^(*n*)^ = *y*^(*n*)^ − *ŷ*^(*n*)^, 1 ≤ *n* ≤ *N*. The mean of the observed data is given by 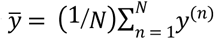. The total sum of squares

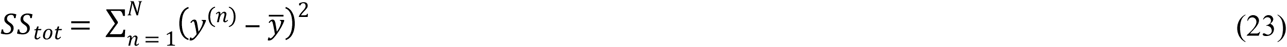

is proportional to the variance of the experimentally observed values. The sum of squares of the residuals is

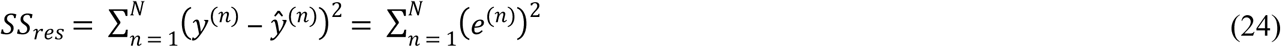

The most general definition of R^2^, as used here, follows from the ratio between these two sums of squares

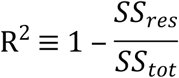

## Acknowledgment

The authors are grateful to prof. Marco Baglietto for the precious discussion about the modelistic part of the work.

## Supporting information

**S1 Fig. Modeling human learning with sequential learning of forward and inverse models.** (a) Temporal evolution of the norm RE of the reaching error as a function of trial number ***n*** calculated from the data of six subjects (black) and from the respective models (blue). (b) Temporal evolution of the norm IME of the inverse model error, the difference between the identity matrix ***I***_***K***_ and the product of the interface map ***H*** and the inverse model ***G***^(***n***)^, estimated from the experimental data (black) and from the model simulations (blue). Both metrics were calculated over a moving window including 12 consecutive trials; the window was shifted forward by one trial at each iteration.

